# Broad bandwidth photoacoustic imaging using a PVDF receiver array

**DOI:** 10.1101/2024.08.20.608753

**Authors:** Sowmiya Chandramoorthi, Antonio López-Marín, Robert Beurskens, Antonius F.W. van der Steen, Gijs van Soest

## Abstract

Photoacoustic (PA) signals are typically broadband, with a correlation between their frequency characteristics and source dimension. The transducers that are commonly used for PA acquisition are optimized for pulse-echo ultrasound imaging and are primarily based on inorganic piezoelectrics in ceramic, single-crystal, or composite form. These transducers are band-limited which limits their functionality as receivers for broadband PA signals. Custom broadband transducers are expensive and complex to manufacture. In this work, we propose to use a poly vinylidene difluoride (PVDF) based transducer for PA acquisition in combination with a commercial single-crystal linear array for pulse-echo acquisition. An 8-element PVDF array with 20dB onboard amplification was built in-house. The PVDF receiver is transparent to the pulse-echo ultrasound, and both transducers were positioned such that they image the same volume. The combined PA raw data from the PVDF and the linear array demonstrated the feasibility of achieving a broader overall reception bandwidth. This study establishes a foundation for a simpler acquisition system that enhances PA signal quality, co-registered with conventional ultrasound imaging, which may support the clinical adoption of PA imaging.

## 1. Introduction

Photoacoustic (PA) imaging is a novel imaging modality that has experienced significant growth and development over the past decade and holds great promise as a diagnostic technology [1-3]. The PA effect entails the generation of an acoustic wave by transient thermoelastic expansion following the absorption of a short light pulse. PA imaging combines ultrasonic resolution and optical absorption contrast to provide functional information in addition to structural information [3]. It has been evaluated in various clinical settings, such as breast cancer staging [4], detection of lymphatics [5], and brain imaging [6]. However, clinical translation of this modality is limited due to the hardware requirements such as high sensitivity and resolution. There is a trade-off between image quality and system complexity; the images from systems that use standard ultrasound (US) echo hardware can be challenging to interpret because of limited sensitivity and reconstruction artifacts, while the acquisition of comprehensive images, that do not exhibit these artifacts, requires highly complex and application-specific imaging systems. The result is a significant threshold for clinical take-up and development of new applications.

PA signals are generally broadband; the received signal frequency content depends on the size distribution of the optical absorbers [7]. For example, in vascular imaging, the vessel sizes range from 0.5 cm to 10 μm, corresponding to frequencies of 300 kHz for the larger vessels to beyond 100 MHz for the microvasculature. Standard US echo probes have a band-limited response, with a typical -6dB fractional bandwidth of up to 80%, relative to the center frequency. As a result, they capture only part of the PA signal, reducing overall sensitivity, particularly to the low and high extremes of the power spectral density function of the PA signal [8]. Due to a combination of PA emission and acoustic attenuation, low frequencies dominate the power spectrum of the PA signal [9], but echo transducers designed for, e.g., vascular, tendon, or nerve imaging lack sensitivity at these low frequencies [10]. The result is that only superficial structures can be imaged effectively.

In recognition of this shortcoming of standard echo hardware for PA imaging, a range of solutions have been proposed in the literature. Manufacturing effective transducers that simultaneously exhibit large enough bandwidth and high sensitivity is not straightforward, since frequency bandwidth is introduced to a mechanically resonant structure by adding damping, which impairs sensitivity. Multi-frequency transducers, using either stacked or laterally offset low- and high-frequency elements, successfully increased bandwidth but often require mechanical scanning to make an image [11-14]. A stacked array concept was also proposed [15]. The application of harmonic ultrasound imaging poses similar challenges [16] and arrays with alternating low- and high-frequency elements have been explored [17]. Some custom-built transducers have been demonstrated for applications such as psoriasis and atherosclerotic plaque [12, 18, 19]. We recently proposed combining imaging data from multiple complementary-frequency US transducers to improve reception bandwidth in photoacoustics [20]. Even though this configuration benefits from targeting a broader spectrum, using two separate linear probes is impractical for clinical settings.

### a. Proposed concept

Piezoceramics and single crystals are commonly used as piezoelectric materials in US transducer fabrication. These materials are well-suited for pulse-echo imaging, credited to their high electromechanical coupling factor. However, they lack bandwidth due to their high acoustic impedance values, typically above 20 MRayl, which create a strongly resonant response. On the other hand, poly vinylidene difluoride (PVDF) is a piezoelectric polymer widely used in the field of sensors and actuators [21]. The low acoustic impedance of PVDF, approximately 4 MRayl, ensures effective coupling with water and soft tissue, which have an impedance of around 1.5 MRayl, contributing to better reception bandwidth than other transducer materials [22, 23]. However, PVDF suffers from lower electromechanical coupling and requires electrical matching. Amplification near the receiving elements (mounted in the holder before the connecting cable) can be used to condition the received signals sufficiently before acquisition by a conventional ultrasound system.

In this work, we propose to use a combination of a custom-designed PVDF receiver, and conventional piezoelectric US linear array transducer, to achieve collocated echo and PA imaging, while capturing most of the PA signal by designing the system to have a broad receive bandwidth. The idea is to mount the acoustically transparent PVDF receiver at the front of a conventional echo probe to add sensitivity for out-of-band signals. In this manner, we improve sensitivity and bandwidth for photoacoustic imaging without adding system complexity or affecting pulse-echo acquisition. In addition, we incorporate the attenuation-dependent sensitivity compensation ratio reported in our previous study [20] to merge data from both probes, that have complementary frequency response. By calibrating the signals from both receivers to an independent standard, we ensure that the signals from multiple complementary US transducers are equalized before combining them, yielding a continuous spectral response and comparable amplitudes in the low and high frequency portions of the signal.

**Fig. 1** shows a schematic of the proposed idea. The acoustically transparent PVDF receiver is mounted on top of a conventional US probe to combine the raw signal data from the imaging region, where the fields of view of the two sensors overlap. Placing contact gel between the linear array, PVDF receiver, and the tissue target ensures acoustic coupling. The signal from the PVDF receiver is amplified by electronics mounted directly adjacent to the sensor; the system is designed to operate at frequencies below the PVDF resonance to maximize bandwidth. Both transducers are connected to an ultrasound data acquisition system to drive the probe and further image processing. The feasibility of the proposed method was tested using simulations and experiments.

**Figure 1:**
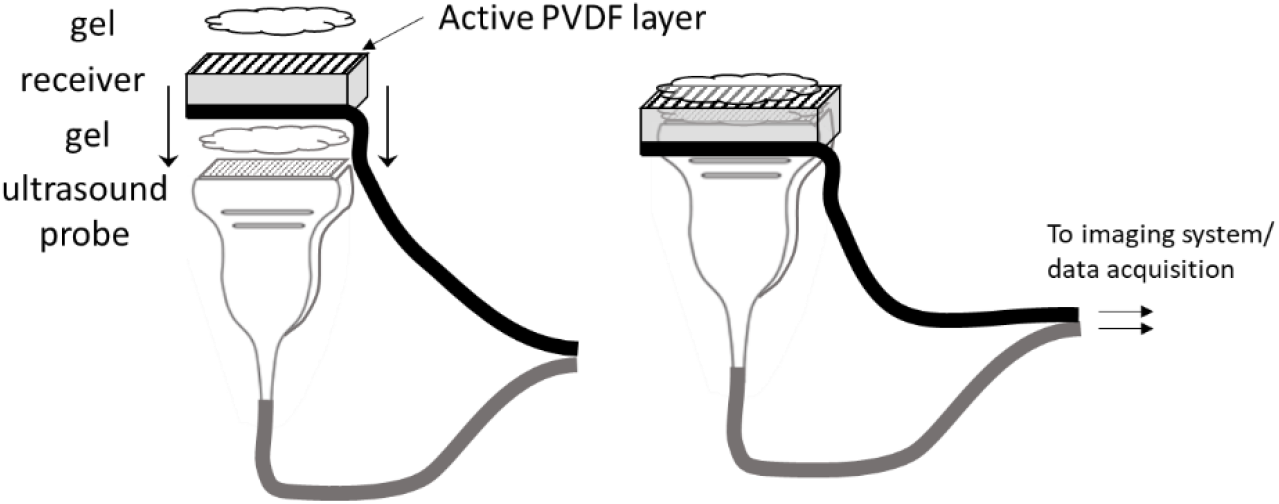
The proposed concept of the broadband receiver: an acoustically transparent piezoelectric material with patterned electrodes can be mounted on a conventional pulse-echo array.

## 2. Methods and materials

### a. Simulations

We analyzed the viability of the design using the finite-element-modeling (FEM) software OnScale (Ansys, Inc., USA). The commercial transducer L22-14vx (Verasonics, Inc., USA) was modeled based on the specifications available from the manufacturer [24]. This probe is a single-crystal 128 element linear array with an 18.5 MHz center frequency and 67% fractional bandwidth. The number of elements in the model was defined as 32 to reduce computation costs and still account for the effect of crosstalk between adjacent elements. **Fig. 2** shows the simulation diagram with the corresponding specifications in Table 1. A PVDF transducer layer and a target were placed 1 mm and 2 mm away from the surface of the probe model, respectively.

**Table 1:** Tabulation of the material specifications.

|  |  |  |
| --- | --- | --- |
| Linear array probe (L22-14vx) | Center frequency | 18.5 MHz |
|  | Material | Single Crystal PMN-28% PT |
|  | Pitch | 0.1 mm |
|  | No of elements | 128 |
| PVDF | Thickness | 52 μm |
|  | Speed of sound | 2250 m/s |
|  | Material | PVDF |
|  | Resonant frequency | 21.63 MHz |
| Backing | Backing, tungsten loaded epoxy, 20% vf, $Z = 8.6 \text{ MRayl}^*$ | |
| Epoxy | Vantico HY1300/CY1301 |  |
| Silicone | Ward Bekker Silicone rubber |  |
| Water | Density | 1000 kgm-3 |
|  | Bulk velocity | 1496 ms-1 |

\* vf - Filament content or fiber volume (% by volume)
Z – Acoustic impedance (MRayl)

**Figure 2:**
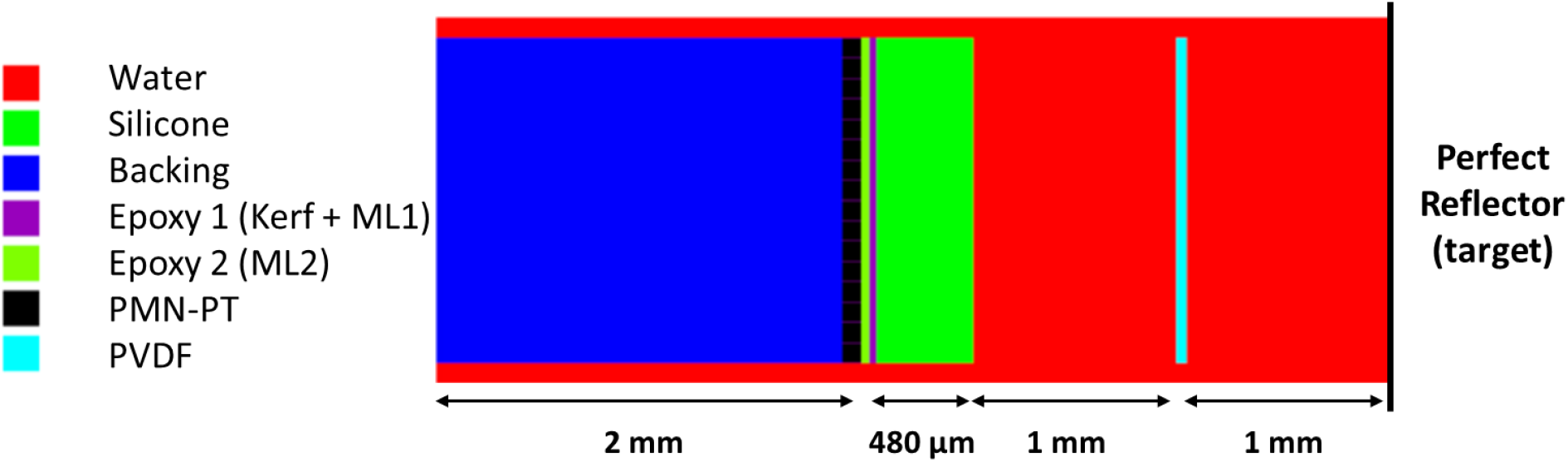
Schematic of the simulation performed on Onscale for the proposed system.

A study was conducted to evaluate the pulse-echo performance of the linear array when altering the thickness of the PVDF transducer layer, including the case when no PVDF was present. In this scenario, a single element close to the center of the probe was driven with a wideband voltage pulse, and the target was configured as an acoustic reflector. The chosen thickness values were 10, 28, 52, and 110 μm, corresponding to commercially available values (Precision Acoustics, Dorchester, UK). The corresponding λ/2 resonance frequencies were calculated as 112.5 MHz, 40.17 MHz, 21.63 MHz, and 10.2 MHz, respectively, using the following equation:

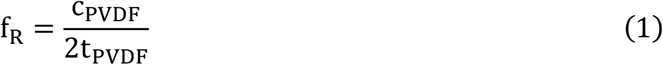

where f_R_, c_PVDF_, and t_PVDF_ are resonance frequency, longitudinal velocity, and thickness of the PVDF layer, respectively.

### b. Experimental setup

A patterned 8-element PVDF array transducer layer was built to test the proposed methodology. The details about the design of the PVDF sheet are tabulated in **Table 2**. The amplifier bandwidth was determined from the frequency response (**Fig. 3a**). The magnitude and phase of the PVDF element impedance were obtained from the spectrum analyzer, shown in **Fig. 3b**.

**Table 2:**
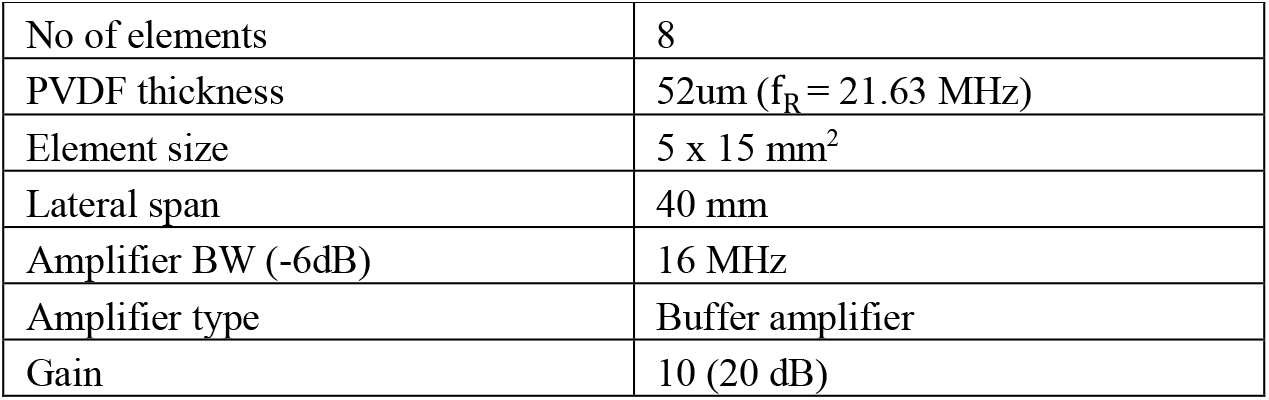
Configuration of the PVDF receiver.

**Figure 3:**
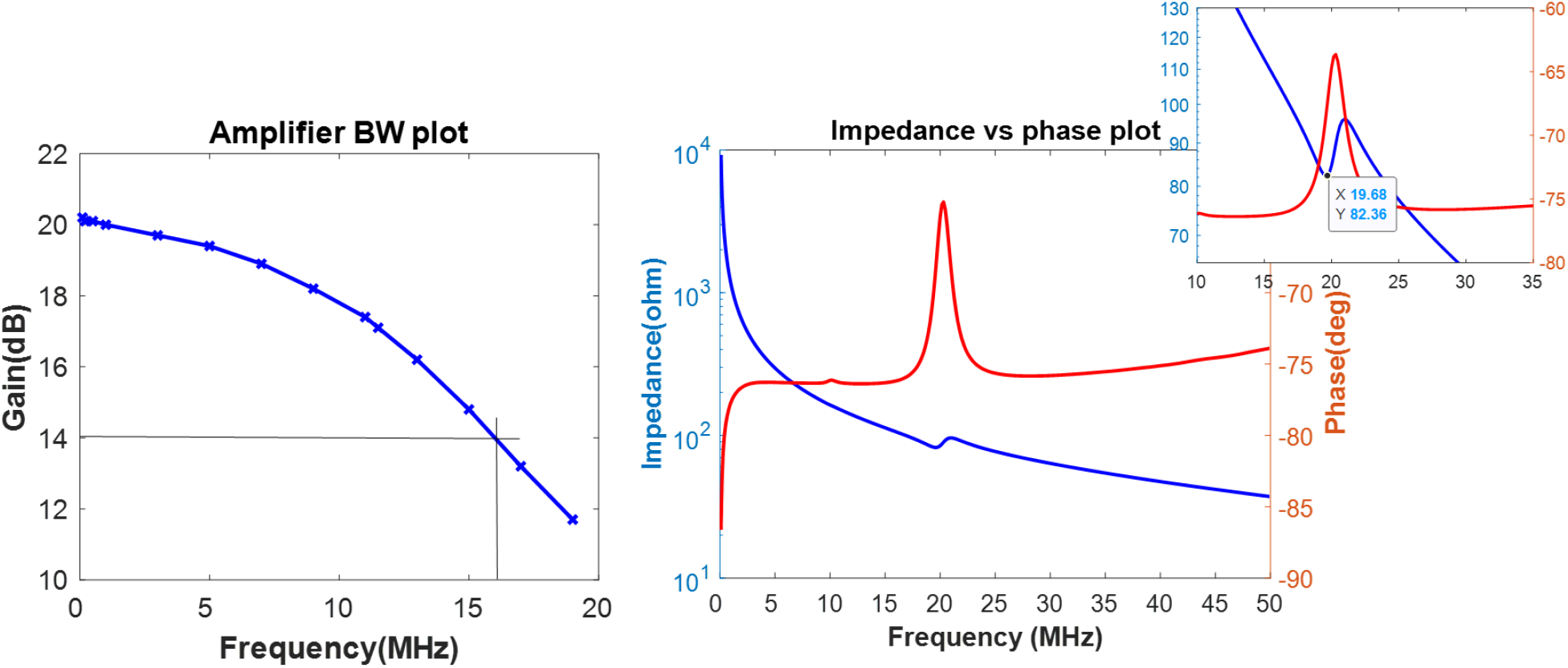
a) Amplifier response as a function of frequency, showing a -6dB bandwidth of nearly 16 MHz; b) impedance (blue curve, left axis) and phase (red curve, right axis) plot of one of the PVDF elements, measured using a spectrum analyser before the amplifier stage

The 8 element PVDF array was built using a 52 μm thick PVDF sheet. The sheet thickness was selected based on its resonant frequency (21.63 MHz) being greater than both the desired receive bandwidth, as well as the pulse-echo frequency. The sheet was patterned into equal parts with each element having a dimension of 0.5 cm by 1.5 cm. The elements were laser patterned using Optec’s nanosecond laser micro-machining system, (Optec S. A, Belgium). The electrical capacitance of each PVDF (∼230 pF) element compared to the capacitance of typical coaxial cables (∼100 pF/m) means that amplification on the ultrasound system side is not optimal [9]. Therefore, we designed an electronic circuit to perform amplification with a drastically lower cable length for this transducer. A 2-layer printed circuit board (PCB) was made with a buffer amplifier (IC ADA4807, Analog Devices, USA) that provides -6 dB bandwidth of 16 MHz and 20 dB gain, see **Fig. 3a**. An acoustic window was cut into the PCB, as shown in **Fig. 4b**, to mount the PVDF and create the closest possible connection to the amplifier circuit.

**Figure 4:**
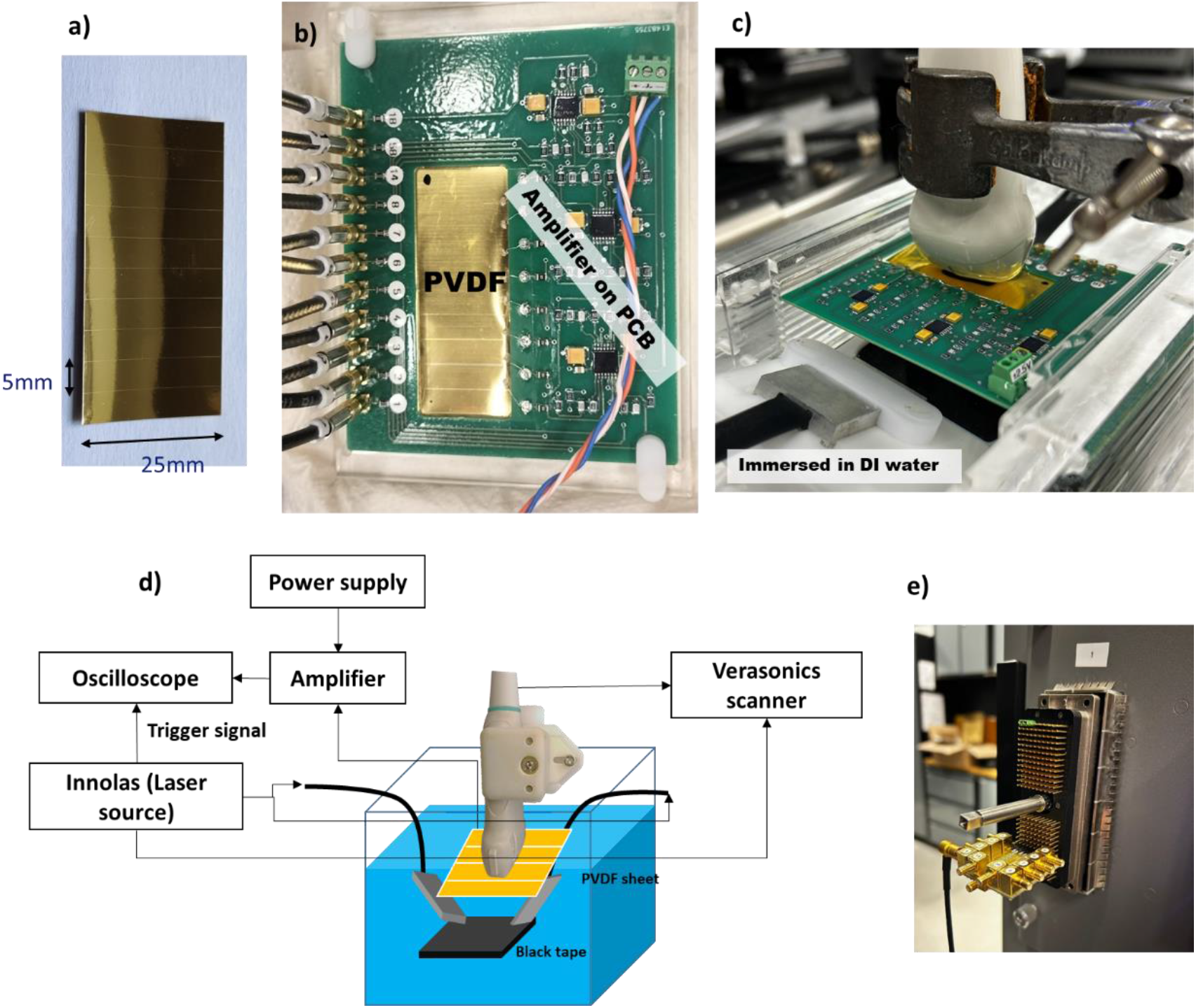
a) The patterned 8 element PVDF array, b) PVDF array after it was mounted on the PCB with onboard amplifier and c) the image acquisition assembly with the linear array positioned on the PVDF and immersed in deionized water. d) Schematic of imaging setup and data acquisition system.

### c. Data acquisition

Combined photoacoustic-ultrasound imaging was performed by immersing the PVDF array and a linear array transducer (L22-14Vx, Verasonics Inc., USA) in deionized water as shown schematically in **Fig. 4d**. An absorbing flat black target and a black nylon wire (300 μm diameter) were illuminated by laser pulses of ∼18 mJ energy, 950 nm wavelength (Spitlight EVO-OPO, Innolas GmbH, Germany). We acquired signals from the PVDF array and the echo probe using a programmable research US system (Vantage 256 HF configuration, Verasonics Inc., USA), both synchronized to the laser light emission (**Fig. 4d**). A custom intermediate PCB was made to interface the PVDF array to one of the UTA-260 S connectors of the Vantage system (**Fig. 4e**). The other UTA connector was used to control the L22-14vx probe. The data acquisition scripts ensured that all receive path parameters such as the TGC, LNA gain, and PGA gain were the same for the PVDF and the echo probe.

In addition, the ground truth photoacoustic response was measured using a 0.2 mm needle hydrophone (Precision Acoustics, UK) from both the PA targets. This data is required to compute the sensitivity compensation ratio, explained in the following section.

### d. Data processing

The raw data acquired from the PVDF elements and the L22-14vx probe were combined to create the broadband signal, using the methodology developed in [20]. Briefly, we equalize the signals from both probes by computing their magnitude relative to the reference provided by the hydrophone standard. The different sensitivity ratios are then used to compensate the signal amplitudes.

The signals acquired on the center element of the PVDF and the L22-14vx probe were averaged over 500 acquisitions. The peak of the photoacoustic signal was windowed from the averaged signal and Fourier transformed to extract the frequency content. The average amplitude of the frequency spectrum recorded by the PVDF and L22-14vx over frequencies within the -3 dB bandwidth was utilized for estimating the sensitivity compensation ratio. The compensation ratios were then obtained by taking a ratio of the amplitudes (PVDF and L22-14vx) in their respective spectral bands, with respect to the ground truth signal recorded by the hydrophone [20].

## 3. Results

### a. Simulation results

We simulated the transmission and reception of a single pulse of 18.5 MHz central frequency from the active piezoelectric layer in the linear array probe through water and the PVDF layer. **Fig. 5a** displays the shape and spectrum of the voltage signal used to drive the piezoelectric layer. The received echo and its corresponding spectrum are plotted in **Fig. 5b**. The plot shows that the designed transducer transmits at a center frequency of 18.5 MHz with a 67% bandwidth in accordance with the specified response of the modeled transducer.

**Figure 5:**
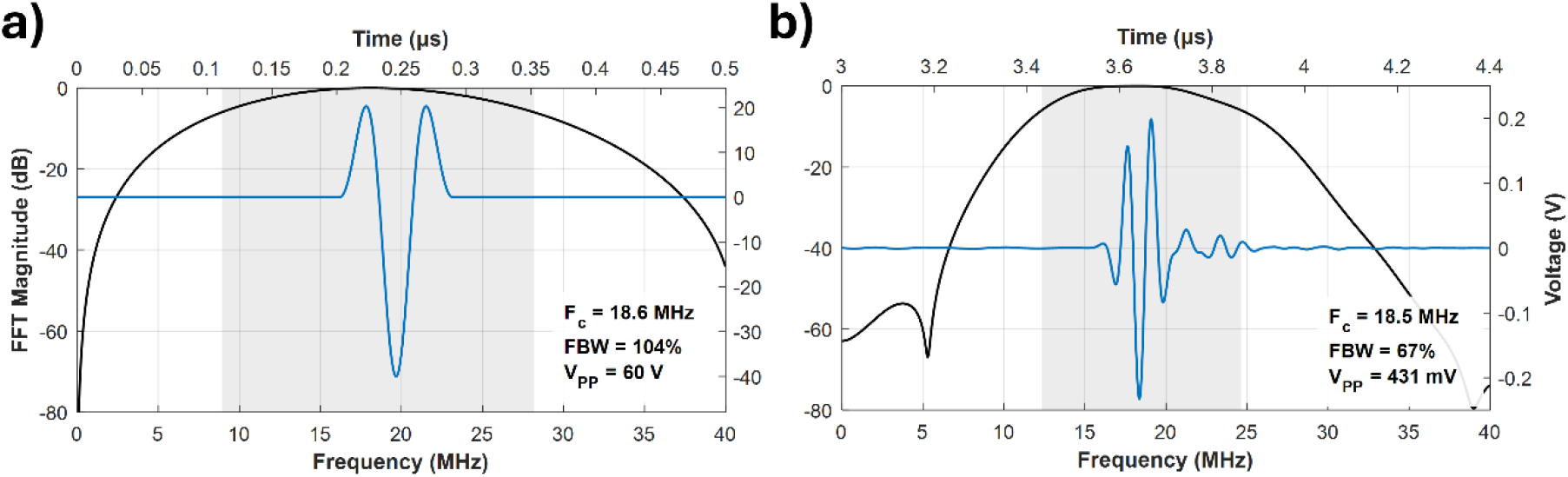
a) Driving signal for the active element in the linear array probe. b) Simulated echo signal and frequency spectrum recorded on the same active element of the linear array. Time signals are represented in blue (right/upper axes), spectra in black and -6 dB fractional bandwidth in grey (left/lower axes). F_c_: central frequency, FBW: -6dB fractional bandwidth, V_PP_: peak-to-peak voltage.

In addition, we analyzed the changes in the pulse-echo signal recorded on the pulse-echo transducer when the intermediate PVDF layer thickness was varied. This analysis was performed in order to understand the impact on pulse-echo signal amplitude and frequency content with the change in PVDF thickness. **Fig. 6(a-d)** shows plots of the echo signals and their corresponding spectra received by the linear array probe element, traveling through PVDF sheets of thickness 10 μm, 28 μm, 52 μm, and 110 μm. The simulation shows a variable effect of the PVDF layer on the pulse-echo signal: for instance, a film of thickness 28 μm exhibits a relatively large attenuation, while the PVDF layer of 110 μm thick has a strong effect on the bandwidth. Notice that the operating band of the pulse-echo probe lies in between the λ/2 resonances of the 52 μm and 110 μm thick layers. The effect of this variation on pulse-echo signal amplitude and bandwidth is shown in **Fig. 5e**. Relevant PVDF thickness values are indicated based on the echo central frequency of 18.5 MHz.

**Figure 6:**
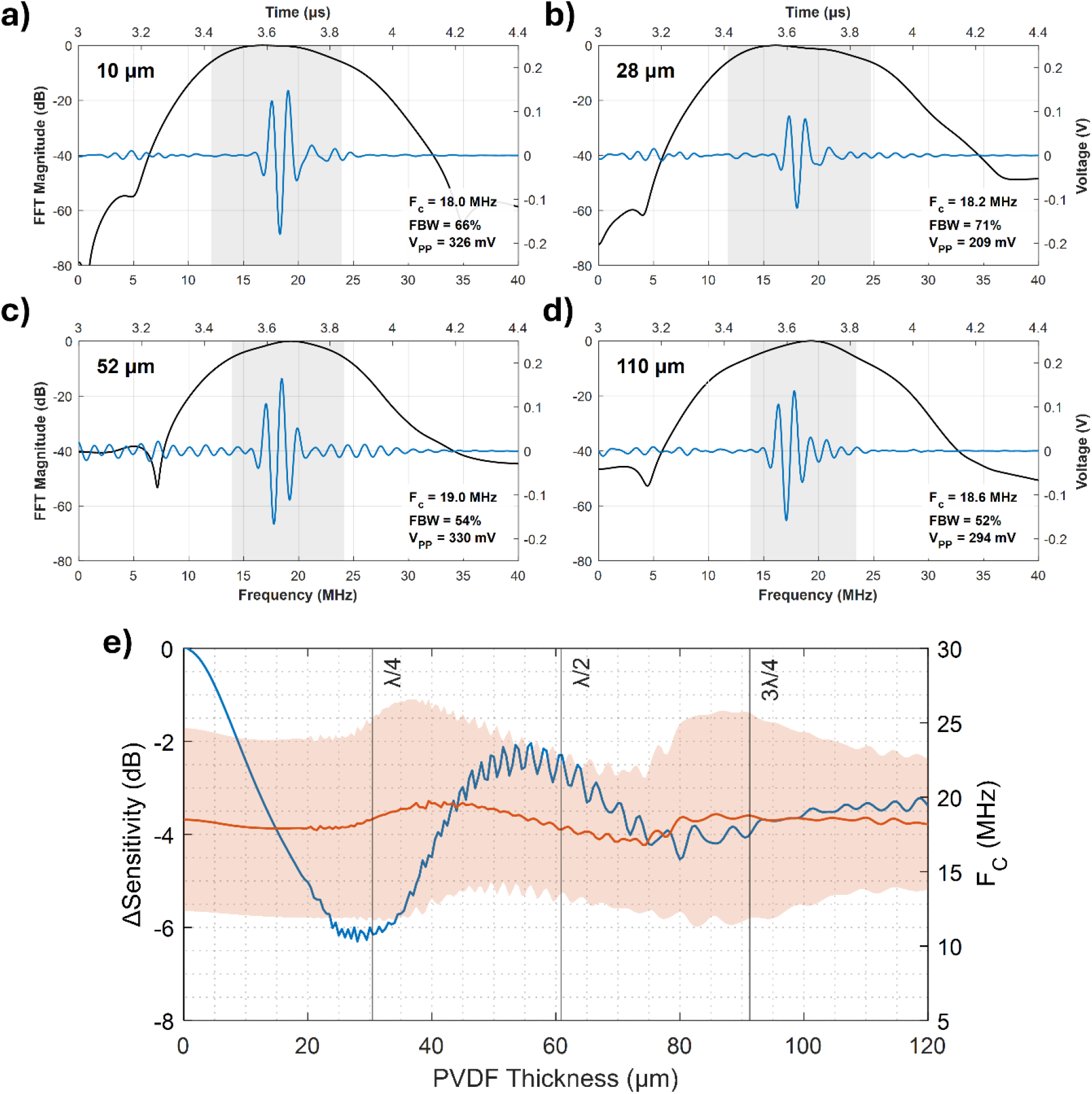
Echo signals recorded on the linear array probe through a PVDF layer of a) 10 μm, b) 28 μm, c) 52 μm and d) 110 μm thickness. Time signals are represented in blue (right/upper axes), spectra in black and -6 dB fractional bandwidth in grey (left/lower axes) e) Sensitivity and bandwidth of the probe measured as a function of PVDF thickness. ΔSensitivity (blue curve) is measured as the ratio of echo peak-to-peak voltage values relative to the zero thickness value. Central frequency (curve) and -6 dB bandwidth (colored area) are represented in red. The λ/4, λ/2, and 3λ/4 thickness values based on a 18.5 MHz working frequency are represented as vertical lines. F_c_: central frequency, FBW: -6dB fractional bandwidth, V_PP_: peak-to-peak voltage.

The performance of the echo signals in pulse-echo mode strongly depends on the thickness of the PVDF sheet. Two acoustic phenomena can alter the received echo: the acoustic impedance mismatch between water and PVDF, which causes reflections to reduce the signal intensity, and the sheet thickness, which affects the contribution of internal reflections within the sheet. A 28-μm sheet, close to a corresponding λ/4 thickness at 18.5 MHz, causes strong attenuation due to destructive interference, minimizing internal reflections and maintaining bandwidth. This effect, though less pronounced, is also observed around 3λ/4 thickness. Conversely, for a 52 μm sheet, close to a λ/2 thickness, the sheet behaves as a resonator producing constructive interference that reduces attenuation but slightly decreases bandwidth. Very thin PVDF sheets cause minimal phase shifts, leading to a lower impact from the internal reflections. Based on these findings, in order to minimize total attenuation in the linear array pulse-echo, we selected commercially available 52 μm PVDF material to build the transducer.

### b. Experimental results

**Fig. 7** shows the photoacoustic signal generated from a broadband target recorded on one of the PVDF array element and its corresponding frequency response, showing a bandwidth of 10 MHz measured at -6 dB level.

**Figure 7:**
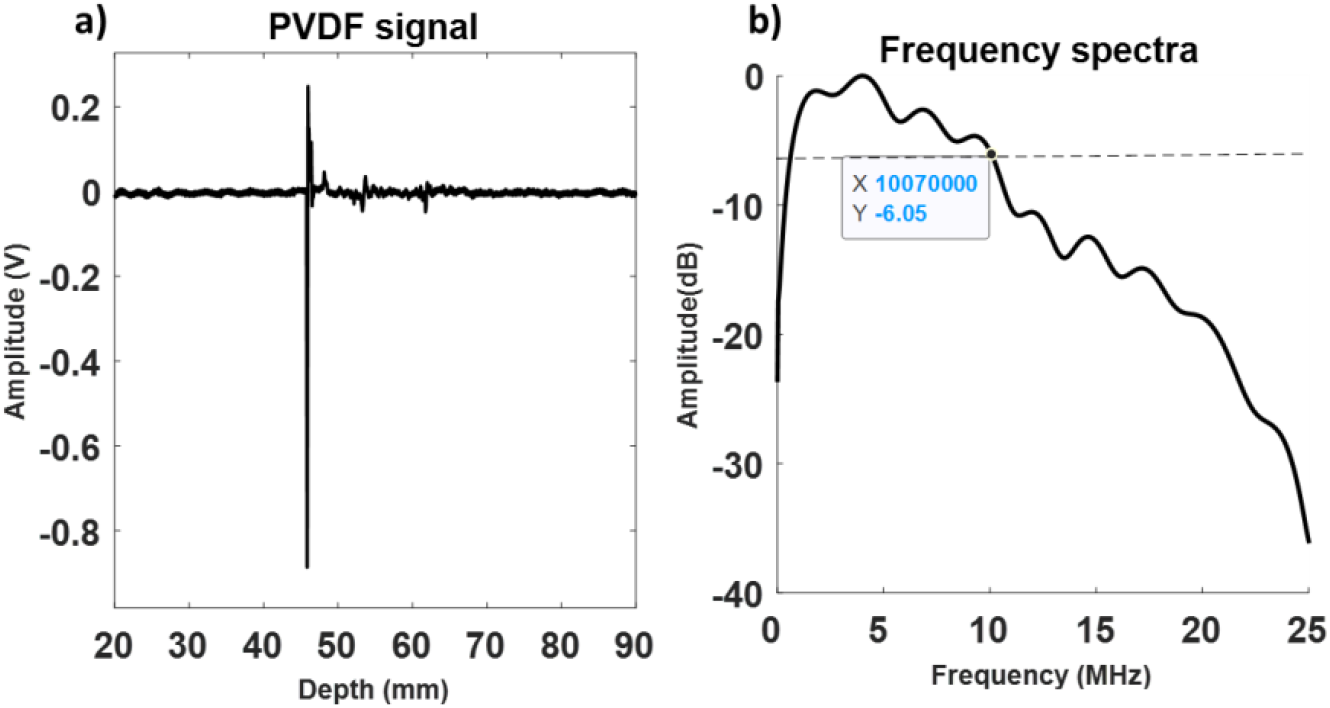
a) Photoacoustic signal recorded on one of the PVDF elements and b) its corresponding frequency spectrum.

**Fig. 8a** and **8d** show the PA signal received by one of the PVDF array elements (blue line), and by the L22-14vx echo probe (red line), from a black broadband target and from a 300 μm nylon wire, respectively. Both signals were equalized and combined at the raw RF level, shown in **Fig. 8b** and **8e**. The frequency spectra of the individual and combined data are plotted in **Fig. 8c** and **8f**, along with the ground truth recorded by a hydrophone from the respective targets. These results demonstrate the complimentary nature of the PVDF and the linear array, which can be usefully deployed for acquiring data over a wider bandwidth, in comparison to the individual arrays.

**Figure 8:**
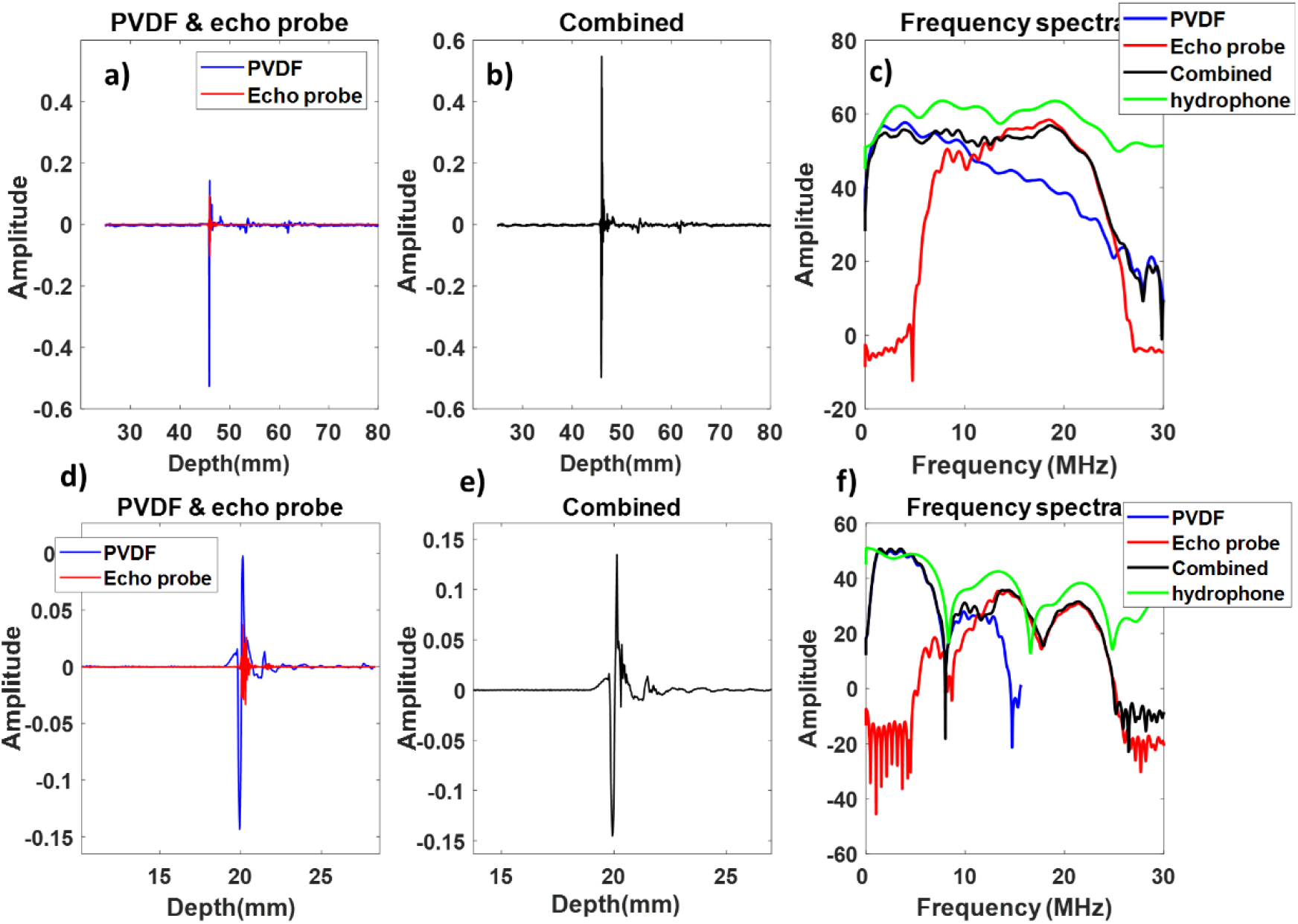
PA signals generated by a black target (a-c) and a 300 μm nylon wire (d-f) recorded on the PVDF (blue lines) and the pulse-echo probe (red lines), respectively. (a, d) Individual signals of the two transducers; (b,e) combined data after equalization; (c, f) corresponding frequency spectra of the individual, combined (black), and hydrophone reference signal (green).

In **Fig. 9** we display experimental results of evaluating the acoustic transparency of the PVDF sheet on pulse-echo and photoacoustic signals recorded on L22-14vx echo probe. **Fig. 9a** and **9b** shows the B-mode images recorded without and with the PVDF. **Fig. 9b** and **9e** show the photoacoustic images recorded on L22-14vx without and with the PVDF. A reflection from the PVDF film is visible in the near field (**Fig. 9b**) which is also marked in **Fig. 9c**. There is also an observable photoacoustic absorption by the film itself that is visible in **Fig. 9e**. A plot of the A line shown in **Fig. 9c** and **9f** demonstrates that the PVDF sheet is relatively transparent and does not affect the pulse-echo and photoacoustic signal amplitude significantly. The attenuation in the pulse-echo signal amplitude with the PVDF present in front of the linear array was 5.03% and the corresponding attenuation in the PA signal was 2.7%.

**Figure 9:**
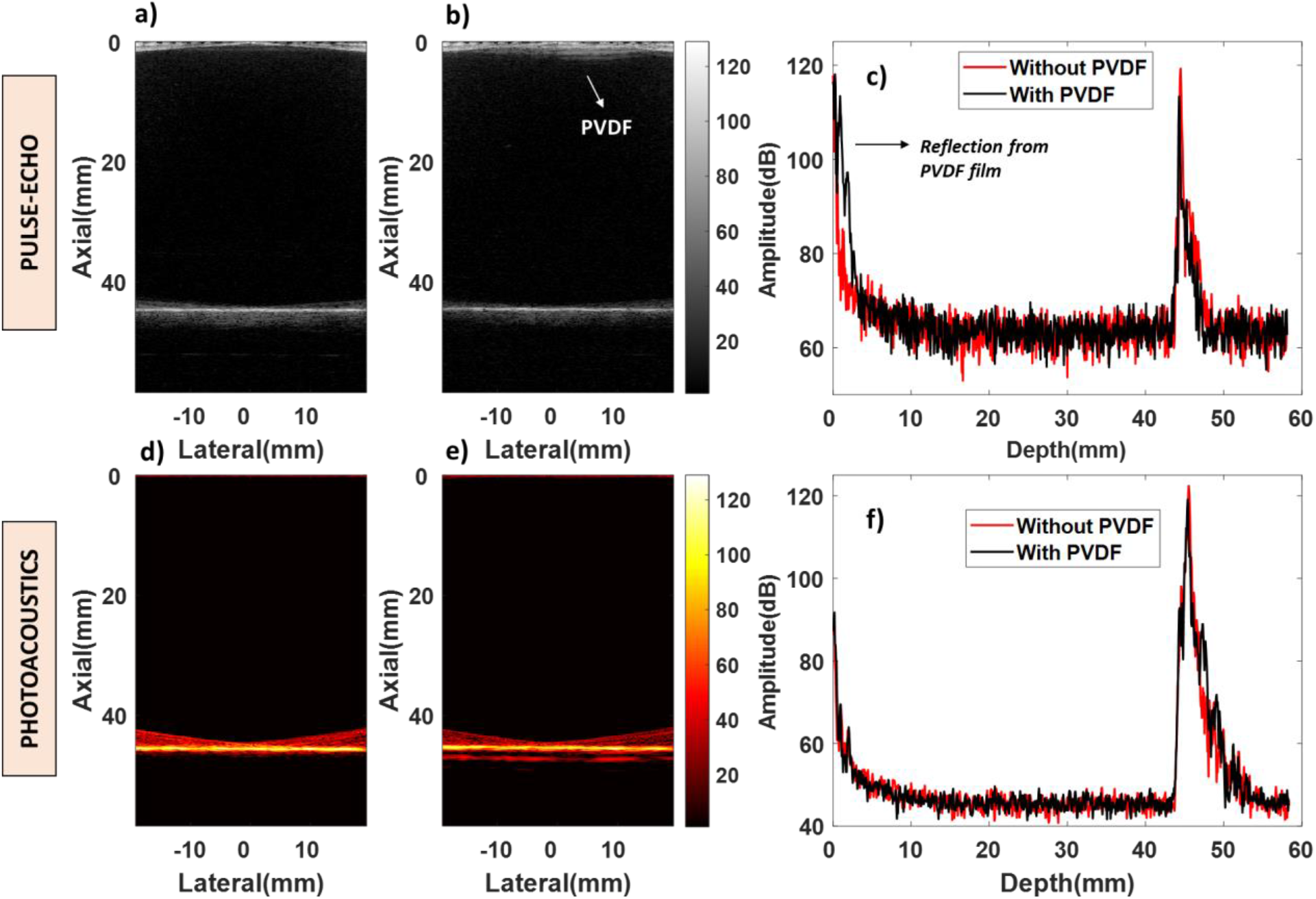
The PVDF sheet is acoustically transparent for pulse-echo as well as for PA signals. Top row: Pulse echo B mode images recorded on the linear array L22-14Vx a) without, and b) with the PVDF film (reflection of the film is visible at the top). c) Comparison of signal amplitudes in RF A-lines without (red) and with (black) PVDF film. Bottom row d)-f) show the corresponding PA signals.

## 4. Summary and discussion

In this work, we have presented a concept for an acoustic receiver that can be used in conjunction with standard echo probes to improve image quality for PA imaging, without the need for customized integrated transducers, which are difficult to manufacture, or special scanners. The goal was to perform PA imaging in a workflow that is similar to echo, using accessible and affordable components. The acoustic receiver was manufactured using PVDF and off-the-shelf electronic components. It was designed to complement a conventional ultrasound array with enhanced sensitivity for low frequencies, where the greatest PA signal power can be expected, thus providing much-needed sensitivity for PA imaging. Its functionality was demonstrated in combination with an L22-14vx transducer, which has a center frequency of 18.5 MHz. We deployed a calibrated method to combine signals from the receiver and the linear array probe and demonstrated that combined PA signals had larger amplitude and bandwidth than those recorded with the linear array alone.

The PVDF-echo probe combination was chosen in such a way that they had complimentary bandwidth. The theoretical bandwidth of the PVDF’s amplifier stretched until 16 MHz, hence the L22-14vx was selected that has a frequency range from 14 to 22 MHz. However, in experiments, the maximum bandwidth that was achievable from the PVDF with a broadband source was ∼10 MHz. This was slightly lower than expected. We hypothesize that this behavior can be attributed to the large surface of the PVDF sheet, which was only approximately flat during the experiment due to its bendable nature. This causes incoherent averaging of the PA signal reaching different regions of the element, which most strongly affects the high frequencies since the path length differences across the element aperture exceed the wavelength, leading to a reduced signal and an apparently smaller bandwidth than expected. Future prototypes will have smaller elements; we will also investigate mounting designs to better maintain the element flatness.

The PVDF array that was manufactured had 8 elements each measuring 15 × 5 mm. This large area leads to a strongly forward-directed sensitivity profile, which means that signals cannot be combined for image reconstruction. In addition, the number of elements was too small to create an image. For this reason, this study was limited to RF lines recorded on each element. These limitations were primarily due to design choices, made to simplify the manufacturing process of this first demonstration prototype.

The results from the simulation study clearly demonstrated the relationship between the PVDF thickness and the effect on the pulse-echo signal recorded on the linear array probe. Ideally, a PVDF layer as thin as possible would be the best choice to reduce the effect on the probe pulse-echo imaging. However, a very thin layer would compromise the receiver’s bandwidth due to the higher amplification requirements. Therefore, a value of 52 μm (close to λ/2 thickness for the center frequency of the echo) was chosen for the PVDF material. In this configuration, the internal reflections within the material are in phase with the echo passing through, contributing to a constructive interference that constrains the attenuation of the transmitted signal, at the cost of reduced bandwidth. The mechanical stability of using a thicker material was also an important consideration during the design of the array, only limited by the resonance frequency of the PVDF sheet, which had to be higher than the desired reception bandwidth. The receiver array presented here has the potential to increase clinical application of PA imaging, by providing an unobtrusive device that enhances image quality compared to standard echo probes. Otherwise, conventional echo hardware can be used, which limits the cost of converting a system to a PA imaging device. The only other component that is needed is the light source, for which affordable, diode-based devices are emerging [25, 26]. Echo hardware and software will need to accommodate two probes simultaneously, a functionality that is becoming available, even on affordable scanners. The receiver thus attempts to address a number of challenges that currently limit real-world use of PA imaging, and we anticipate it may accelerate the take-up of the technology in clinical diagnostics.

## CRediT author statement

Sowmiya Chandramoorthi: Conceptualization, Methodology, Investigation, Formal analysis, Writing – original draft.

Antonio López-Marín: Conceptualization, Methodology, Investigation, Writing - review & editing.

Robert Beurskens: Methodology, Writing - review & editing.

Antonius Van der Steen: Writing – review & editing, Supervision.

Gijs Van Soest: Conceptualization, Methodology, Writing – review & editing, Supervision.

## Data availability

The authors do not have permission to share data.

## Acknowledgments

This work was supported by the Dutch Research Council NWO (Vici-16131).

## References

1. Wang, L.V., Photoacoustic Imaging and Spectroscopy. 1st Edition ed. 2009, Boca Raton: CRC Press.

2. Beard, P., Biomedical photoacoustic imaging. Interface Focus, 2011. 1(4): p. 602–31.

3. Riksen, J.J.M., A.V. Nikolaev, and G. van Soest, Photoacoustic imaging on its way toward clinical utility: a tutorial review focusing on practical application in medicine. J Biomed Opt, 2023. 28(12): p. 121205.

4. Neuschler, E.I., et al., A Pivotal Study of Optoacoustic Imaging to Diagnose Benign and Malignant Breast Masses: A New Evaluation Tool for Radiologists. Radiology, 2018. 287(2): p. 398–412.

5. Van Heumen, S., et al., LED-based photoacoustic imaging for preoperative visualization of lymphatic vessels in patients with secondary limb lymphedema. Photoacoustics, 2023. 29(2213-5979 (Print)): p. 100446.

6. Na, S., et al., Massively parallel functional photoacoustic computed tomography of the human brain. Nat Biomed Eng, 2022. 6(5): p. 584–592.

7. Diebold, G.J., T. Sun, and M.I. Khan, Photoacoustic monopole radiation in one, two, and three dimensions. Phys Rev Lett, 1991. 67(24): p. 3384–3387.

8. Ku, G., et al., Multiple-bandwidth photoacoustic tomography. Phys Med Biol, 2004. 49(7): p. 1329–38.

9. Daeichin, V., et al., A Broadband Polyvinylidene Difluoride-Based Hydrophone with Integrated Readout Circuit for Intravascular Photoacoustic Imaging. Ultrasound Med Biol, 2016. 42(5): p. 1239–43.

10. Szabo, T.L. and P.A. Lewin, Ultrasound transducer selection in clinical imaging practice. J Ultrasound Med, 2013. 32(4): p. 573–82.

11. Kalkhoran, M.A., F. Varray, and D. Vray, Dual frequency band annular probe for volumetric pulse-echo optoacoustic imaging. Proceedings of the 2015 Icu International Congress on Ultrasonics, 2015. 70: p. 1104–1108.

12. Luo, X., et al., Stack-Layer Dual-Element Ultrasonic Transducer for Broadband Functional Photoacoustic Tomography. Front Bioeng Biotechnol, 2021. 9: p. 786376.

13. Sibo, L., et al. A dual-layer micromachined PMN-PT 1–3 composite transducer for broadband ultrasound imaging. in 2013 IEEE International Ultrasonics Symposium (IUS). 2013.

14. Liu, J.H., et al., Design, fabrication and testing of a dual-band photoacoustic transducer. Ultrasonic Imaging, 2008. 30(4): p. 217–227.

15. Wada, T., Photoacoustic imaging apparatus, photoacoustic imaging method, and probe for photoacoustic imaging apparatus. 2013, FUJIFILM CORPORATION (Tokyo, JP): United States.

16. Martin, K.H., et al., Dual-Frequency Piezoelectric Transducers for Contrast Enhanced Ultrasound Imaging. Sensors, 2014. 14(11): p. 20825–20842.

17. Neer, P.L.M.J.V., et al., Super-harmonic imaging: development of an interleaved phased-array transducer. IEEE Transactions on Ultrasonics, Ferroelectrics, and Frequency Control, 2010. 57(2): p. 455–468.

18. Cao, Y., et al., Highly sensitive lipid detection and localization in atherosclerotic plaque with a dual-frequency intravascular photoacoustic/ultrasound catheter. Transl Biophotonics, 2020. 2(3): p. e202000004.

19. Aguirre, J., et al., Precision assessment of label-free psoriasis biomarkers with ultrabroadband optoacoustic mesoscopy. Nature Biomedical Engineering, 2017. 1(5).

20. Chandramoorthi, S., et al., Wideband photoacoustic imaging in vivo with complementary frequency conventional ultrasound transducers. Frontiers in Physics, 2022. 10.

21. Ting, Y., et al., Design and characterization of one-layer PVDF thin film for a 3D force sensor. Sensors and Actuators a-Physical, 2016. 250: p. 129–137.

22. Xiao, J., et al., Photoacoustic endoscopy with hollow structured lens-focused polyvinylidine fluoride transducer. Appl Opt, 2016. 55(9): p. 2301–5.

23. Sappati, K.K. and S. Bhadra, Piezoelectric Polymer and Paper Substrates: A Review. Sensors (Basel), 2018. 18(11).

24. Verasonics. L22-14v High Frequency Imaging Probe Datasheet. 2017 2017–08; Available from: https://verasonics.com/resources-downloads/.

25. Kohl, A., et al., An ultra compact laser diode source for integration in a handheld point-of-care photoacoustic scanner, in Photons Plus Ultrasound: Imaging and Sensing 2016. 2016.

26. Kuniyil Ajith Singh, M. and W. Xia, Portable and Affordable Light Source-Based Photoacoustic Tomography. Sensors (Basel), 2020. 20(21).

